# Heat tolerance, oxidative stress response tuning, and robust gene activation in early-stage *Drosophila melanogaster* embryos

**DOI:** 10.1101/2024.04.29.591747

**Authors:** Emily E. Mikucki, Thomas S. O’Leary, Brent L. Lockwood

## Abstract

In organisms with complex life cycles, life stages that are most susceptible to environmental stress may determine species persistence in the face of climate change. Early embryos of *Drosophila melanogaster* are particularly sensitive to acute heat stress, yet tropical embryos have higher heat tolerance than temperate embryos, suggesting adaptive variation in embryonic heat tolerance. We compared transcriptomic responses to heat stress among tropical and temperate embryos to elucidate the gene regulatory basis of divergence in embryonic heat tolerance. The transcriptomes of tropical and temperate embryos were differentiated by the expression of relatively few genes, including genes involved in oxidative stress. But most of the transcriptomic response to heat stress was shared among all embryos. Further, embryos shifted the expression of thousands of genes and showed robust gene activation, demonstrating that, contrary to previous reports, early embryos are not transcriptionally silent. The involvement of oxidative stress genes in embryonic heat tolerance corroborates recent reports on the critical role of redox homeostasis in coordinating developmental transitions. By characterizing adaptive variation in the transcriptomic basis of embryonic heat tolerance, this study is a novel contribution to the literature on developmental physiology and genetics, which often lacks ecological and evolutionary context.

## INTRODUCTION

All life is inherently sensitive to sudden changes in temperature due to the disruption of the weak bonds that stabilize macromolecular structures (Somero et al., 2017). Survival against sudden, acute increases in temperature, such as would be encountered during a heat wave or fever, necessitates cellular coping strategies, like the heat shock response, that aid cells in mitigating macromolecular damage (Ashburner and Bonnert, 1979; Feder and Hofmann, 1999; Lindquist, 1986; Ritossa, 1962; Tomanek, 2010). The heat shock response is a fast transcriptional response, characterized by the differential regulation of thousands of genes (Buckley et al., 2006; Gasch et al., 2000; Leemans et al., 2000), that confers whole-organism heat tolerance (Feder et al., 1996; Welte et al., 1993). Consequently, variation in transcriptional regulation in response to heat stress has been the target of thermal selection in many species (Bettencourt et al., 2002; Liu et al., 2017; Lockwood et al., 2010; Whitehead et al., 2011). Without the heat shock response, organisms die from mild heat stress (Hofmann et al., 2000), and thus cellular responses to acute heat stress are critical to surviving climate warming, as the global climate crisis is leading to increases in mean temperatures and also the frequency of heat waves (Marx et al., 2021; Perkins-Kirkpatrick and Lewis, 2020).

The reliance on transcriptional regulation for heat stress tolerance poses physiological constraints that may limit adaptive responses to heat waves. Any organism that lacks a robust heat shock response is ill equipped to survive a heat wave (Hofmann et al., 2000), and any population that lacks variation among individuals in the heat shock response lacks the ability to adapt to future heat waves (Kelly et al., 2012). Thus, physiological constraints to coping with heat stress may influence biogeographic range limits and ultimately determine species persistence in a warming world (Somero, 2020; Tomanek, 2010). In species with complex life cycles, the life stage with the greatest degree of sensitivity to heat stress will most likely set its thermal limits (Coyne et al., 1983; Lockwood et al., 2018). In the context of the heat shock response, the earliest stages of embryogenesis may be particularly thermally sensitive, as early embryos in all animals and endosperms in higher plants lack a fully developed transcriptional machinery (Baroux et al., 2008; Tadros and Lipshitz, 2009). But despite the potential limits to heat tolerance in early life stages, previous work has demonstrated genetic variation in heat tolerance in early life stages that correlates with variation in the thermal environment (Feiner et al., 2018; Lockwood et al., 2018; Sato et al., 2015), suggesting that thermal adaptation is possible even when there appear to be physiological constraints to coping with heat stress in early life stages.

Because early life stages are often thermally sensitive, they are likely to be critical to the thermal ecology of many species (Angilletta et al., 2013). Yet, early life stages are rarely studied in ecologically relevant thermal contexts (Feiner et al., 2018; Gleason et al., 2018; Lockwood et al., 2018). Embryos of *Drosophila melanogaster* are immobile, and thus at the mercy of the environment where their mothers lay them. Within hours, the temperature of a piece of necrotic fruit, the preferred oviposition site for *D. melanogaster*, may increase by as much as 20°C (Roberts and Feder, 2000). Unlike adult flies, embryos cannot avoid thermal extremes through behavioral thermoregulation (Allemand and David, 1976; Dillon et al., 2009; Huey and Pascual, 2009). Moreover, early-stage *D. melanogaster* embryos are more thermally sensitive than all other life stages (Gilchrist et al., 1997; Lockwood et al., 2018; Tucic, 1979; Welte et al., 1993). Previous studies have suggested that the thermal sensitivity of early embryos stems from their reduced transcriptional activity (Graziosi et al., 1980; Lockwood et al., 2017; Welte et al., 1993). However, the extent to which maternal provisioning of mRNAs may protect early embryos against heat stress has not been fully explored (but see Lockwood et al., 2017). Further, natural variation in the transcriptome of early embryos of *D. melanogaster* has not been previously measured, and transcriptomic responses to acute heat stress in early embryos have never been characterized.

Here we characterized the transcriptomes of tropical and temperate *D. melanogaster* early embryos in the context of acute heat stress, and hereby elucidate molecular physiological mechanisms that underlie adaptive thermal tolerance. *D. melanogaster* has a broad global distribution (Kapun et al., 2021), and previously we demonstrated that tropical *D. melanogaster*, collected from relatively hot locations around the globe, produce embryos that are more heat tolerant than embryos of temperate flies collected from relatively cool locations in North America (Lockwood et al., 2018). Fundamentally, the physiology of early embryogenesis is controlled by gene activation (Ali-Murthy et al., 2013; Fowlkes et al., 2008; Wieschaus, 1996), and thus we sought to characterize transcriptomic responses to elucidate the potential mechanisms that underlie this adaptive divergence.

Embryonic heat tolerance could manifest in several ways at the level of the transcriptome. On a mechanistic level, tropical embryos may have higher heat tolerance because they express key genes involved in heat stress coping mechanisms, such as genes that encode molecular chaperones (Feder et al., 1996; Lockwood et al., 2017; Richter et al., 2010; Tomanek and Somero, 2000). Alternatively, tropical embryos may be more transcriptionally active across their whole transcriptome due to an earlier onset of gene activation. Our results allow us to address these possibilities to determine how the transcriptome mediates responses to environmental variability in a developmental context. Moreover, to our knowledge this is the first study to compare the early-life transcriptomes of broadly sampled *D. melanogaster* populations around the globe. Thus, we present an account of the transcriptomic responses that not only distinguish heat tolerant and heat sensitive genotypes, but that also represent a large extent of the transcriptomic diversity of the *D. melanogaster* species.

## MATERIALS AND METHODS

### Fly strains

We propagated five tropical and five temperate North American iso-female lines of *D. melanogaster* as described previously (Lockwood et al., 2018). Notably, these ten lines exhibit continuous variation in embryonic heat tolerance, as indexed by survival (LT_50_) following acute heat stress (45 min), with the tropical lines having the highest heat tolerance (Fig. 1). All tropical lines were obtained from the Drosophila Species Stock Center with origins from Mumbai, India (MB) (Stock number: 14021-0231.45); Accra, Ghana (GH) (Stock number: 14021-0231.182); Monkey Hill, St. Kitts (SK) (Stock number: 14021-0231.34); Chiapas de Corzo, Chiapas, Mexico (CP) (Stock number: 14021-0231.22); and Guam, USA (GM) (Stock number: 14021-0231.198). All Vermont lines were collected from East Calais, VT (Cooper et al., 2014). Iso-female lines were established by taking single females whose progeny were inbred for several generations to isogenize the genetic background within each line. This is a well-established technique to minimize the potential for lab evolution by removing genetic variability (Cooper et al., 2014). We maintained flies at controlled densities of 50-100 adults per vial (95mm x 25mm) under common-garden conditions in incubators (DR-36VL, Percival Scientific Inc., Perry, Iowa, USA) set to 25°C, 55% relative humidity, and a 12L:12D light cycle on a cornmeal-yeast-molasses medium for at least 2 generations prior to collecting embryos for embryonic heat shock. 25°C represents an ecologically relevant temperature that both tropical and North American *D. melanogaster* flies are exposed to in the wild.

**Figure 1.**
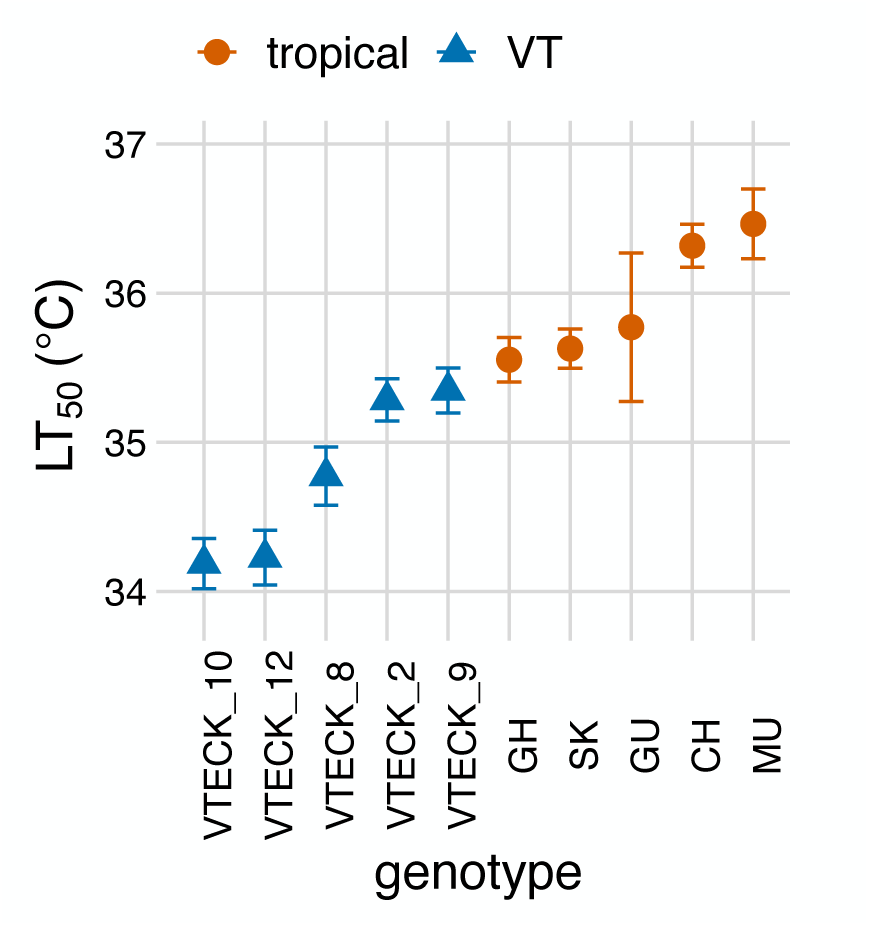
*D. melanogaster* embryonic heat tolerance as indicated by LT_50_—the temperature that causes 50% mortality following a 45-min. exposure—among the ten genotypes used in this study. For details see (Lockwood et al., 2018).

### Embryonic heat shock

Heat shocks mimicked the natural, acute increases in temperature of the necrotic fruit on which *D. melanogaster* oviposit their eggs (Roberts and Feder, 2000). Further, heat shock temperatures spanned the range of LT_50_ values of the ten isofemale lines (Fig. 1), which mimicked the design used in the original study that characterized the LT_50_ phenotype (Lockwood et al., 2018). We allowed three- to five-day-old adult flies (n ≈ 100 pairs) from each of the 10 lines to lay eggs on grape juice agar plates (60 x 15 mm) dabbed with active baker’s yeast for one hour at 25°C. We then wrapped the egg plates with parafilm, and submerged them in a circulating water bath (A24B, Thermo Scientific, Waltham, MA, USA) for 45-minutes at one of four temperatures: 25°C, 32°C, 34°C, and 36°C. In the water bath the temperature of the plates increased at a rate of +0.4°C^-1min^ until reaching the target temperature (Lockwood et al., 2018). We repeated the heat shocks for a minimum of three replicates per treatment (genotype x temperature).

After the heat shock, we rinsed the eggs from each plate with 1x embryo wash solution (0.12 M NaCl, 0.05% (v/v) Triton X-100), for approximately 30 seconds in order to remove residual yeast. We then dechorionated the embryos by washing with a 50:50 solution of water and bleach for one minute, followed by a rinse in dH_2_O for 30 seconds. We removed residual water by blotting with a Kimwipe. We pooled eggs (n ≥ 30) from each plate, placed them into 1.5 mL microcentrifuge tubes, and immediately flash froze them in liquid nitrogen. Pools of embryos were preserved at −80°C until further transcriptomic analysis.

### RNA extraction and sequencing

We extracted total RNA by homogenizing the pooled embryos in 250 µL of TRIzol (Tri-reagent, Sigma Life Science, St. Louis, MO, USA), and centrifuging all samples at 4°C at 6,000g for one minute (Sorvall ST89, Thermo Scientific). We then added an additional 150 µL of trizol (Tri-reagent, Sigma Life Science) and 150 µL of chloroform (Sigma Life Science) to each sample, followed by centrifugation at 4°C at 12,000 x g for ten minutes (Sorvall ST89, Thermo Scientific). Lastly, we precipitated each sample using isopropanol and ethanol. To determine the quantity of RNA in each extraction, we used a Nanodrop spectrophotometer (NanoDrop 2000, Thermo Scientific). To assess the quality of RNA, we used agarose (2%) gel-electrophoresis (EL-200, Walter, Plymouth, MI, USA) to visualize bands of RNA. We only sequenced samples that contained >1 µg of RNA and that exhibited strong bands, corresponding to intact 28S and 18S ribosomal RNAs.

We sent all samples (n = 120) to Novogene (Sacramento, CA, USA) for 150 base-pair paired-end mRNA sequencing. Further quality control of each RNA sample was performed by Novogene using a Nanodrop spectrophotometer, agarose gel electrophoresis, and the Agilent 2100 BioAnalyzer (Santa Clara, CA, USA). Only 110 of the 120 samples passed this second round of quality control; thus, we report results from 110 samples. The samples excluded were as follows: BO at 34°C, BO at 36°C, CP at 36°C, GH at 32°C, GM at 32°C, VT12 at 25°C, VT2 at 32°C, VT2 at 36°C (x2), and VT9 at 25°C.

After quality control, libraries were prepped by enriching the mRNA using oligo(dT) beads, then cDNA was synthesized using mRNA templates and random hexamer primers. Double-stranded cDNA libraries were sequenced using the Novaseq 6000 (S4 Flowcell) (Illumina, San Diego, CA, USA). Library quality was assessed using Qubit 2.0 Fluorometer (Thermo Scientific), Agilent 2100 BioAnalzyzer, and q-PCR.

We checked the quality of paired-end raw sequence reads using FastQC (v. 0.11.5) (Andrews, 2010). We trimmed the forward and reverse reads using Trimmomatic (v. 0.36) (Bolger et al., 2014) to remove adapter sequences and low quality leading and trailing bases by cropping the first 12 bases of each read, and the 20 leading and trailing bases. We also scanned the reads with a 6-base sliding window, cutting the read when the average quality per base was below a Phred score of 20. Lastly, we dropped reads below lengths of 35 bases.

### Quantification, normalization, and statistical comparisons of transcript abundances

We used salmon (v. 1.1.0) to map transcripts to the *Drosophila melanogaster* transcriptome (FlyBase v. r6.34) and to quantify the abundance of transcripts for each of the 110 embryonic samples (Patro et al., 2017). We used R (v. 4.2.3) (R Core Team, 2023) and the package “DESeq2” (v. 1.32.0) to normalize read counts and perform statistical analysis (Love et al., 2014). There were 27,303 transcripts with non-zero total read count that were sequenced and mapped to the *D. melanogaster* transcriptome. To determine transcripts with significantly different abundances due to region of origin (tropical or temperate North America) and temperature, we used the likelihood ratio test (LRT) within the DESeq function. From these tests, we generated lists of differentially abundant transcripts based on an FDR < 0.05 for the two factors individually, and the interaction between heat shock temperature and region. 8,081 and 5,974 transcripts were deemed by DESeq2 to have low counts in the region and temperature contrasts, respectively.

We conducted a principal component analysis to describe the major axes of variation in transcript abundances among samples using the prcomp function in R. To convert transcripts to genes, we used the R package gprofiler2 (Kolberg et al., 2020).

### Coexpression analysis

To identify modules of co-expressed transcripts that may be correlated with LT_50_ or temperature, we performed weighted gene co-expression network analysis (WGCNA) with the wgcna package in R (Langfelder and Horvath, 2008). For this analysis, following the recommendation of the authors of WGCNA, we filtered out transcripts with fewer than 15 counts in more than 75% of samples. This rendered 8,754 transcripts, which we used to construct the network. We performed variance stabilization to normalize the counts of the filtered dataset with the vst function in the DESeq2 package. We then constructed a signed network with a soft-power threshold of 10, which provided a good model fit (R^2^ = 0.91; Fig. S1) while maintaining a high degree of connectivity (mean connectivity = 161.6). We used a merge cut height of 0.25, which merged modules that were 75% similar in their clusters of transcripts. These parameters produced 17 modules of co-expressed transcripts (Fig. 4A).

### Functional enrichment analysis

We conducted functional enrichment analysis with the R package VISEAGO (Brionne et al., 2019). We focused our analyses on biological process categories and ran the classic algorithm of overrepresentation analysis using the Fisher’s exact test with an FDR cut-off of 0.01. The broad functional categories shown in Figure S4B are not GO terms and were curated by the authors to present a broad classification of the cellular and developmental processes in the study system. We used FlyBase (release FB2023_06) (Jenkins et al., 2022), along with the published literature, to classify gene function.

## RESULTS

### Transcriptomic correlates of embryonic heat tolerance

Overall, embryonic heat tolerance (Fig. 1) was associated with the expression of relatively few genes, as embryonic transcriptomes were largely similar regardless of region of origin (Fig. S2). 793 genes showed differences in expression between tropical and VT embryos (DESeq2, likelihood ratio test, region factor, FDR <0.05), with 285 of these genes also showing significant changes in expression following heat stress (Fig. S3). These 793 genes were encoded by 799 mRNA transcripts, and most of these were more highly expressed in tropical embryos than VT embryos—429 transcripts were more abundant in tropical vs. 370 transcripts that were more abundant in VT embryos. Overall, 3,826 genes encoded by 4,135 transcripts showed significant changes in expression following heat stress (Fig. S3; DESeq2, likelihood ratio test, temperature factor, FDR<0.05), indicating a robust heat shock response, but this response was largely shared among all embryos (see below).

There was little functional overlap between the genes that were differentially expressed between tropical and VT embryos and the heat stress response genes (Fig. 2A; Table S1). 72 GO function categories distinguished tropical and VT embryonic transcriptomes (Fig. 2A), and nine of these were associated with the oxidative stress response (Fig. 2B). Of note, many GO categories associated with cellular stress were enriched among the genes that changed expression upon heat stress, but only the oxidative stress response was different between tropical and VT embryos (Fig. 2B). Among the 17 genes in the “response to oxidative stress” GO category, eight were more highly expressed in tropical embryos and seven were more highly expressed in VT embryos, with two genes having two transcript isoforms each that were more highly expressed in either tropical or VT embryos (Fig. 3). The specific functional roles of these genes could be classified as contributing to either the prevention of the formation of reactive oxygen species (ROS) or resistance to oxidative stress (Table 1). Notably, four of the 17 genes were involved in ROS prevention, and three of these were more highly expressed in tropical embryos (Table 1; Fig. 3).

**Figure 2.**
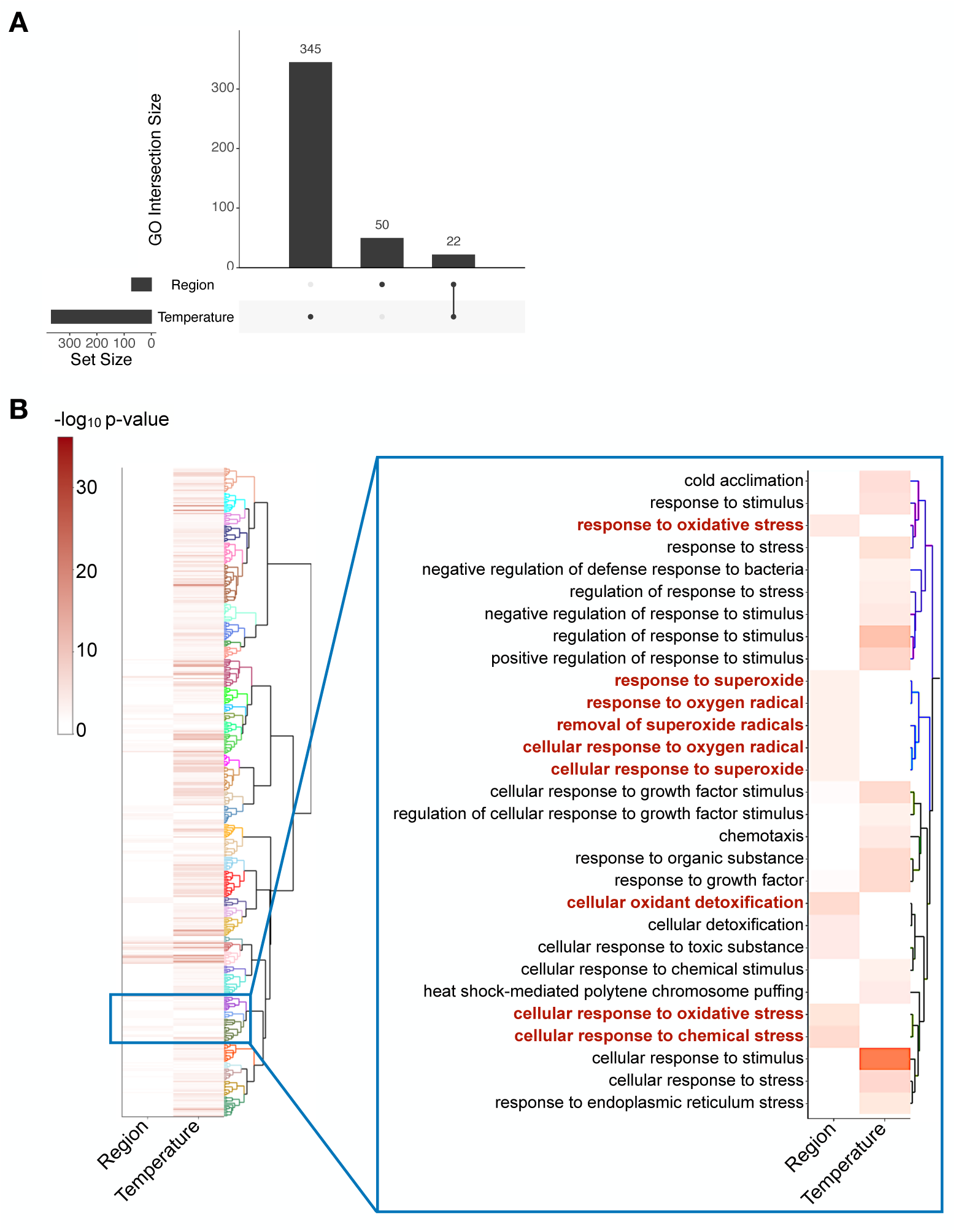
Functional enrichment of GO biological process categories among the genes significantly differentially expressed between embryos of different regions (tropical vs. temperate) and/or across temperature (DESeq2, LRT, FDR < 0.05). (A) The number of enriched GO categories in each group of genes and overlap between groups. (B) Heat map depicting the degree to which GO categories were enriched, with the darker color indicating greater enrichment. The dendrogram indicates the relationships among GO categories, with functionally related categories clustered together on the tree. The inset highlights stress response and response to stimulus GO categories, with oxidative stress response categories highlighted in red.

**Figure 3.**
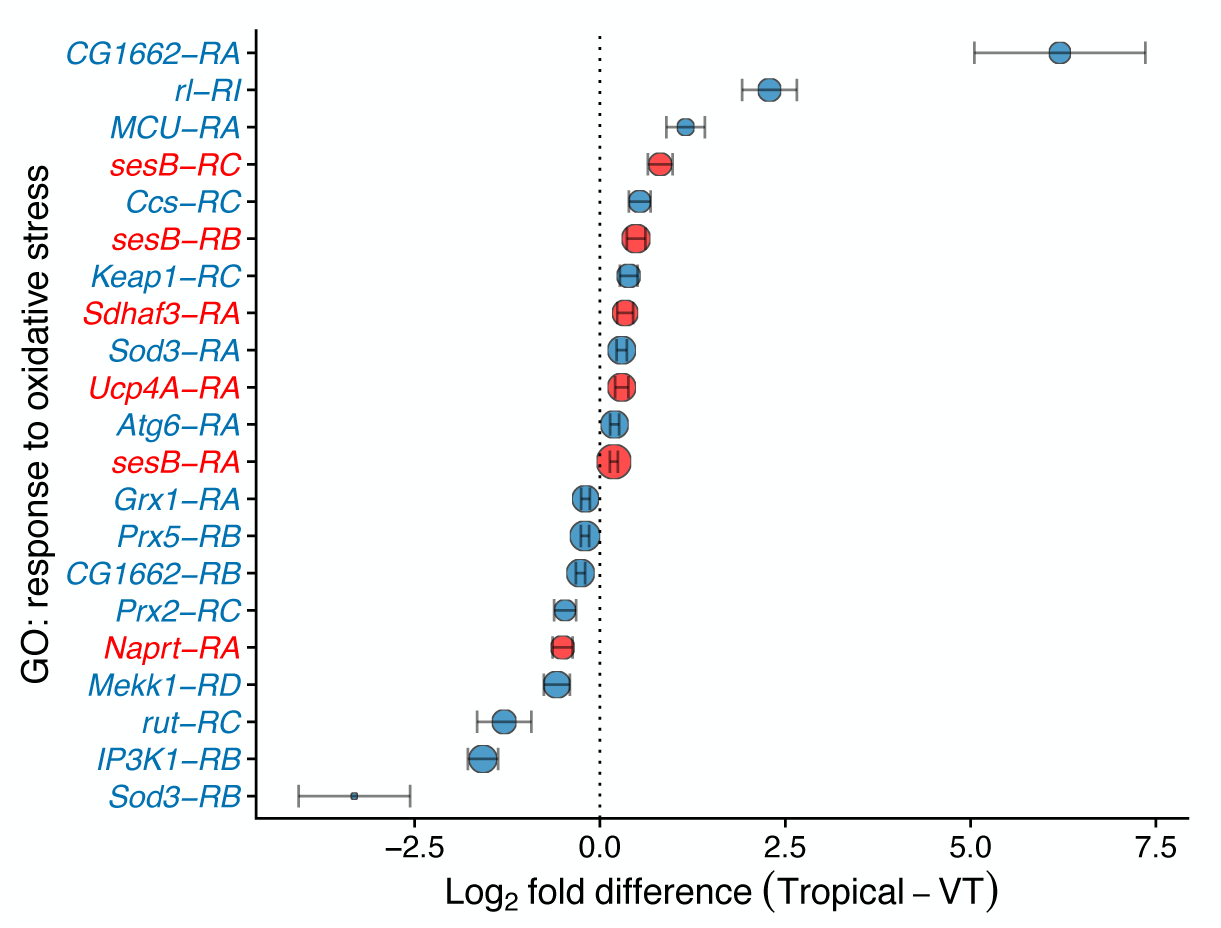
Mean differences in expression of oxidative stress response transcripts between tropical and Vermont embryos across all temperatures. Positive numbers indicate higher expression in tropical embryos. Negative numbers indicate higher expression in Vermont embryos. Transcripts in blue are involved in general resistance to oxidative stress, either by direct scavenging or a cellular response to ROS. Transcripts in red are involved in prevention of ROS formation. The size of the points indicates mean normalized expression (Log_2_ scale). Error bars indicate standard error of the mean.

**Table 1.** Relative expression of genes involved in the oxidative stress response (GO:0006979) and their functional roles.

| Gene | Higher expressed | Role | Mechanism |
| --- | --- | --- | --- |
| *sesB* | tropical | prevention | maintenance of redox state via ATP transport |
| *Sdhaf3* | tropical | prevention | chaperone of SdhB (Complex II of ETC) |
| *Ucp4A* | tropical | prevention | maintenance of redox state via uncoupling of ETC and ATP synthetase |
| *rl* | tropical | resistance | MAPK signaling stress response |
| *MCU* | tropical | resistance | regulation of oxidative stress induced apoptosis |
| *Ccs* | tropical | resistance | direct scavenging via chaperoning of Sod1 |
| *Keap1* | tropical | resistance | regulation of oxidative stress induced gene expression |
| *Atg6* | tropical | resistance | regulation of autophagy in response to stress |
| *Naprt* | VT | prevention | maintenance of redox state via NAD+ synthesis |
| *Grx1* | VT | resistance | direct scavenging of ROS via glutathione pathway |
| *Prx5* | VT | resistance | direct scavenging of ROS |
| *Jafrac1* | VT | resistance | direct scavenging of ROS |
| *Mekk1* | VT | resistance | direct scavenging of ROS via regulation of dual oxidase |
| *rut* | VT | resistance | cAMP signaling stress response |
| *IP3K1* | VT | resistance | calcium signaling stress response in the ER |
| *CG1662* | mixed | resistance | direct scavenging of ROS via peroxisome |
| *Sod3* | mixed | resistance | direct scavenging of ROS |

Using weighted gene correlation network analysis (WGCNA)(Langfelder and Horvath, 2008), we identified six modules of gene expression, representing a total of 1,617 genes, whose expression was correlated with variation in LT_50_ (Figs. 1 and 4; Table S2). Notably, these modules were not only correlated with LT_50_, but also distinguished transcriptional responses between tropical and VT embryos, in both overall RNA abundance levels and the response to acute heat stress (Figs. 4B-G). Note that we refer to module names based on the color labels that were assigned by the WGCNA analysis.

**Figure 4.**
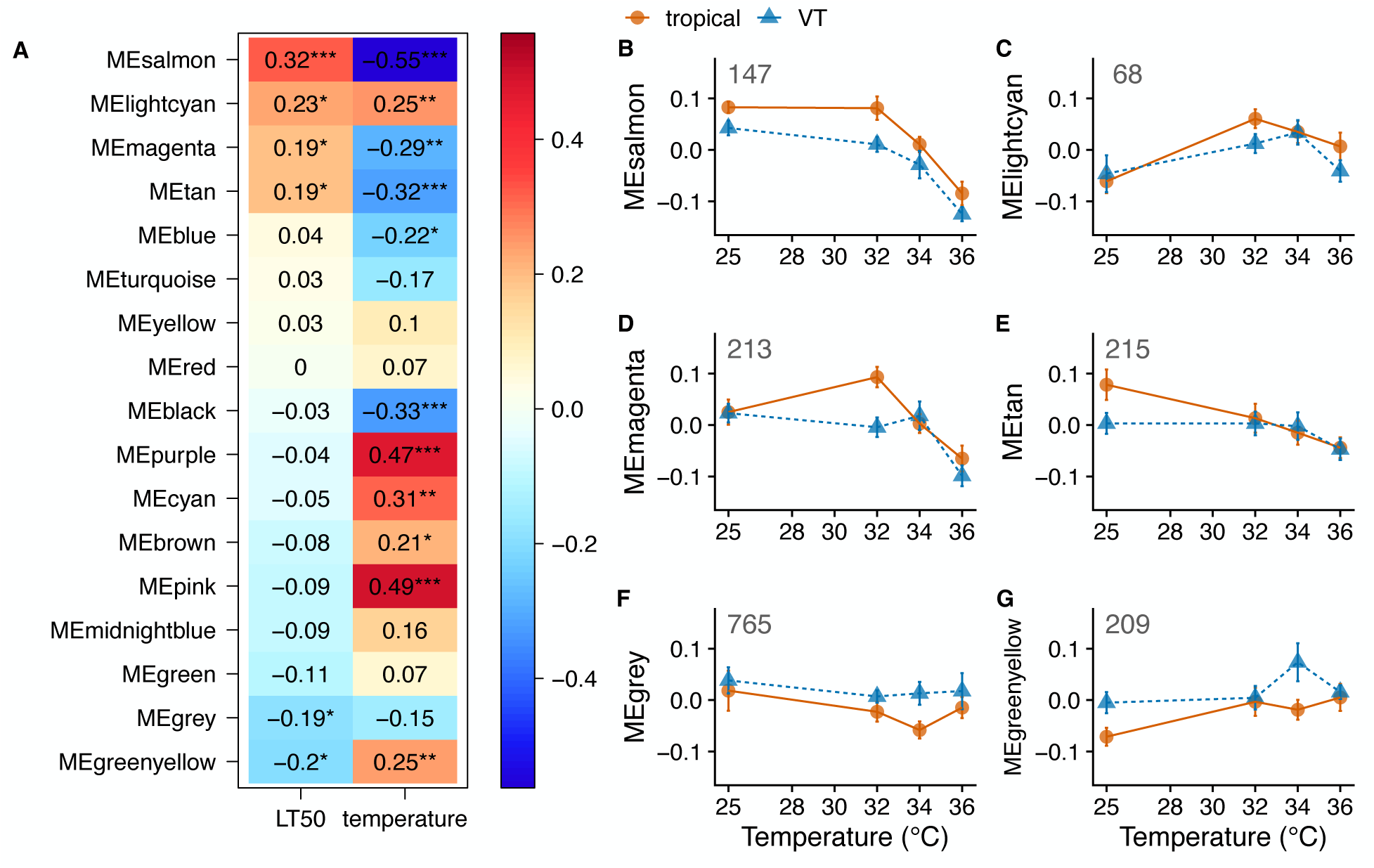
Modules of gene expression identified by WGCNA. (A) Pearson correlation coefficients between module eigengenes (ME) and LT_50_ or temperature. Red colors indicate positive correlations, and blue colors indicate negative correlations. Asterisks indicate level of significance of the Pearson correlation: * *P*<0.05, \*\**P*<0.01, \*\*\**P*<0.001. (B-G) ME values across temperature for the six modules with significant correlations to LT_50_. Note that ME reflects the pattern of expression common to all genes in a particular module, with higher values indicating greater abundance of RNA. Numbers in each panel indicate the number of genes in that module.

Most modules that were significantly correlated with heat tolerance had module eigengene values that were positively correlated with LT_50_ (Fig. 4A), and thus contained genes whose expression was higher in embryos with higher heat tolerance. For example, the salmon module had module eigengene values that were higher in tropical embryos across all temperatures (Fig. 4B), whereas the light cyan and magenta modules showed higher module eigengene values in tropical embryos at 32°C (Figs. 4C and 4D). Overall, the gene expression patterns most correlated with higher embryonic heat tolerance were characterized by higher expression in tropical embryos, either at baseline (25°C) or at the lowest heat shock temperature (32°C), suggesting that preparation prior to heat shock and the initial response to heat shock were transcriptomic signatures of higher embryonic heat tolerance.

On the other hand, two modules had module eigengene values that were negatively correlated with LT_50_, and thus contained genes whose expression was higher in VT than tropical embryos (Figs. 4A, 4F, and 4G). These two modules highlight differential responses between tropical and VT embryos to 34°C; the grey module had genes that were down regulated at 34°C only in tropical embryos, whereas the green-yellow module had genes that were upregulated at 34°C in VT embryos.

Functional enrichment analysis of these six WGCNA modules revealed a diverse molecular signature of embryonic heat tolerance (Fig. S4). Overall, each WGCNA module consisted of genes involved in GO categories that were distinct from other modules (Fig. S4A). However, GO categories could be grouped into higher-order functional families that were shared among multiple modules (Fig. S4B). While the salmon and magenta modules included many genes spread across a wide array of functional categories, the light cyan module consisted of genes in only certain categories, including RNA processing, transcription, protein and lipid metabolism, and regulatory pathways. Similarly, the tan module had a relatively specific GO signature, with genes involved in protein and lipid metabolism, morphogenesis, localization and transport, and organellar organization. The two modules negatively correlated with heat tolerance, grey and green-yellow, had a predominance of genes involved in RNA processing, transcription, and protein and lipid metabolism. Interestingly, the green-yellow module had many genes involved in histone modification, indicating a higher-order mechanism for differential transcriptional regulation between heat tolerant and heat sensitive embryos.

### The embryonic transcriptomic response to heat stress

As mentioned above, the transcriptomes of both tropical and temperate embryos were responsive to temperature, with thousands of transcripts changing abundance following heat stress (Fig. 5). Most of the transcripts (2,921 transcripts) were downregulated following heat stress, while 1,214 transcripts were induced (Fig. 5). We did not detect any genes for which there were region-specific responses to temperature (DESeq2; LRT, region x temperature interaction). This was despite identifying WGCNA modules with region-specific temperature responses (as reported above). We interpret this discrepancy to be the result of (1) false-discovery rate correction, which is implicit in the gene-by-gene statistical testing implemented by DESeq2, but which is absent from the network-based approach of WGCNA, and (2) the dynamic nature of expression changes across temperature, for which the DESeq2 likelihood ratio test is less likely to identify as significant interactions because the direction and effect sizes of the interactions changed at different temperatures. Nonetheless, we include the results of both sets of analyses because we believe that they each highlight important aspects of the data.

**Figure 5.**
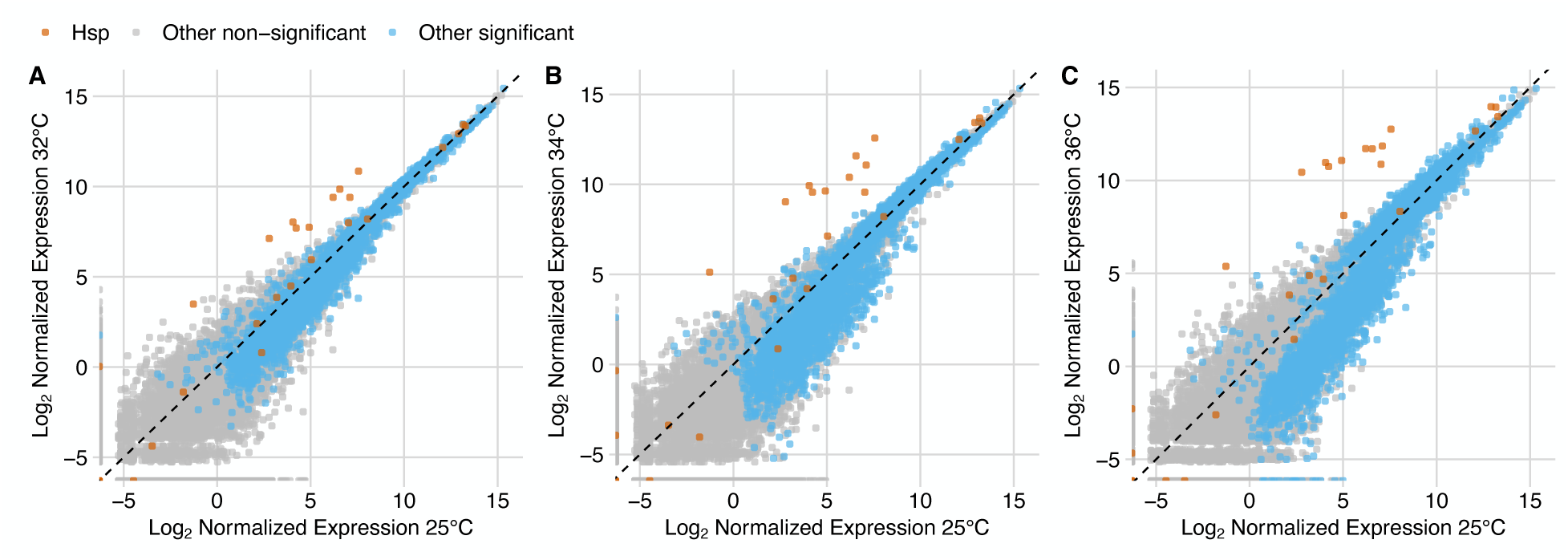
The shared embryonic heat shock response. Plotted are mean log_2_-normalized expression at 25°C vs. 32°C (A), 34°C, (B), or 36°C (C). Red dots are heat shock protein genes (Hsp), gray dots are non-significant non-heat-shock-protein genes, and blue dots are significant non-heat-shock-protein genes (FDR < 0.05). Diagonal dashed line is the 1:1 line of the x and y axes.

Based on WGCNA clustering, we identified six modules, representing 2,772 genes, with eigengenes that were significantly correlated to temperature and not heat tolerance (Figs. 4A and S5). Thus, these modules represent the transcriptional response to heat stress that all embryos shared and which followed a bimodal pattern, with some modules decreasing and others increasing in expression. GO analysis revealed a diverse array of functional categories that were highly enriched in these modules (Table S3). Notably, the modules that decreased in expression (i.e., blue and black) contained genes involved in translation (GO:0006412) and peptide biosynthesis (GO:0043043). Whereas the modules that increased in expression (i.e., purple, cyan, brown, and pink) contained genes involved in cellular localization (GO:0051641), cellular component organization (GO:0016043), and the cellular response to stress (GO:0033554). Molecular chaperone activity was evident in the pink module, with protein refolding (GO:0042026), chaperone-mediated protein folding (GO:0061077), and heat-shock mediated polytene chromosome puffing (GO:0035080) being among the top 5 most enriched categories (Table S3). Indeed, the pink module contained many of the classic heat shock genes (e.g., *Hsp70 Bb*, *Hsp68*, and *Hsp23*), all of which were highly induced following heat stress (Figs. 5 and S5).

## DISCUSSION

### Expression of few genes underlies embryonic heat tolerance

Taken together, our data suggest that most of the embryonic transcriptome is conserved and that the differential regulation of a relatively small fraction of the transcriptome confers heat tolerance. *Drosophila* embryogenesis has been shown to be unperturbed across a range of more benign temperatures, from 18°C to 29°C (Houchmandzadeh et al., 2002). Thus, it seems that a potential explanation for the pattern of overall transcriptional homogeneity across heat shock temperatures and among embryos from different geographic regions may be the highly conserved nature of early embryonic development (Davidson, 1986), such that developmental processes have evolved to be robust to environmental and mutational perturbations—i.e., embryonic transcriptomes are canalized (von Heckel et al., 2016; Waddington, 1942). Indeed, gene expression in early development is highly evolutionarily conserved within the *D. melanogaster* species, as well as across the *Drosophila* genus, particularly among maternally loaded transcripts (Omura and Lott, 2020). But despite this high degree of conservation, thermal adaptation of embryonic heat tolerance was not constrained, as tropical and temperate embryos maintain a measurable difference in their heat tolerances (Fig. 1) (Lockwood et al., 2018). Previous studies in other species have shown that overall transcriptomic responses to heat stress are conserved, but differences in the expression of a few key genes underlie heat tolerance (Dilly et al., 2012; Lockwood et al., 2010). In adults of *D. melanogaster* and the closely related species *D. simulans*, populations from Maine and Panama exhibited transcriptomic divergence in responses to more benign temperature exposures of 21°C and 29°C (Zhao et al., 2015). But, here too, most genes were not differentially expressed between populations. Our data demonstrate that *Drosophila* embryos are consistent with this overall pattern.

An alternative explanation for the lack of variation among transcriptomes is that thermal selection acts on levels of biological organization other than the transcriptome. There is evidence supporting this possibility in the form of the evolution of thermal stability of proteins (Dong et al., 2018; Fields and Somero, 1998; Gu and Hilser, 2009), which may influence the temperature at which the cellular stress response is induced (Tomanek and Somero, 2000). But the process of protein evolution depends on the relatively slow accumulation of amino-acid-changing mutations (Chin et al., 2022), such that adaptive divergence across the proteome likely occurs after speciation events. Thus, the short-term evolutionary timescales that separate divergent populations within a species may be too short to render this as a plausible explanation for the data we report on herein. Nonetheless, we cannot rule out the possibility of thermal adaptation via other levels of biological organization that we did not characterize. But we also emphasize the likelihood that thermal selection acts on the transcriptome, as demonstrated in fish, lizards, *Daphnia*, and copepods (Campbell-Staton et al., 2021; Feiner et al., 2018; Herrmann et al., 2018; Schoville et al., 2012; Whitehead et al., 2011). Given the out-sized role of gene activation in coordinating early developmental events (Li et al., 2014; Wieschaus, 1996), we believe that transcriptional regulation is likely to be critical to any developmental phenotype, including embryonic heat tolerance.

In terms of physiological mechanisms, genes involved in the oxidative stress response appear to be central to embryonic heat tolerance. This was one of the most consistent patterns that we observed for any functional category of genes—i.e., tropical and Vermont embryos exhibited differential regulation of a diverse set of oxidative stress genes. Some oxidative stress genes encode proteins that actively scavenge ROS after they are formed, such as *Sod3* (Fridovich, 1998; Jenkins et al., 2022), while other oxidative stress genes encode proteins that prevent ROS from forming altogether, such as *Ucp4A* (Echtay, 2007; Jenkins et al., 2022; Somero et al., 2017). Both tropical and Vermont embryos expressed oxidative stress genes involved in scavenging of ROS or otherwise supporting stress resistance. However, tropical embryos were distinguished in their ability to have higher expression of genes involved in the prevention of ROS, such as *Ucp4A*, *sesB*, and *Sdhaf3*. Because tropical embryos are more heat tolerant and exhibited these patterns of gene expression, the prevention of oxidative stress may be a key physiological mechanism underlying thermal adaptation in early *Drosophila* embryos.

The link between thermal stress tolerance and oxidative stress tolerance has been observed across a broad range of taxa (Abele et al., 2002; De Zoysa et al., 2009; Teixeira et al., 2013; Tomanek and Zuzow, 2010), which corroborates these observations. Thermal stress can lead to oxidative stress via increased production of free radicals and other ROS leading to changes in cellular redox potential (Kültz, 2005; Somero et al., 2017). While ROS can cause cellular damage by oxidizing DNA, RNA, lipids, and/or proteins (Gleason et al., 2017; Schieber and Chandel, 2014), ROS are also important cellular signaling molecules that are involved in the regulation of a vast array of homeostatic cellular processes via control of cellular redox state (Schieber and Chandel, 2014; Sies, 2017; Sies and Jones, 2020). Indeed, recent studies have highlighted the critical role of cellular redox state for the coordination of key developmental transitions (Petrova et al., 2018; Syal et al., 2020), cell fate determination (Syal et al., 2020), and energy metabolism (Hocaoglu and Sieber, 2022; Hocaoglu et al., 2021) during embryogenesis. Because ROS are key to normal cellular and developmental function, antioxidant systems are important for maintaining the delicate balance of ROS to ensure the maintenance of cellular redox state (Sies and Jones, 2020). An excess of ROS can lead to redox imbalance that results in cellular dysfunction via disruption of metabolism and cell signaling (Schieber and Chandel, 2014). Altogether, our results suggest that heat stress produces excess ROS that disrupt embryonic development and that more heat tolerant embryos are better able to prevent ROS formation and thus maintain redox homeostasis during heat stress.

Beyond the oxidative stress response, a wide array of cellular and developmental processes appears to be involved in producing embryos with higher heat tolerances. Indeed, the expression of hundreds of genes was correlated with LT_50_, including genes involved in RNA processing, histone modification, morphogenesis, and ion homeostasis. So, even though a relatively small fraction of genes differentiated the transcriptomes of tropical and Vermont embryos, this small fraction performed a diverse set of functions. Because of the pervasive effects of temperature (Scholander, 1955; Somero et al., 2017) it is not surprising that a diverse set of genes was associated with heat tolerance. In addition, these genes exhibited a specific pattern of expression in response to increasing levels of heat stress. More heat tolerant embryos tended to have higher expression of these genes prior to heat stress or upon experiencing low levels of heat stress. Previous studies have also demonstrated that more heat tolerant organisms tend to exhibit a more robust early transcriptional response to acute heat stress than more heat sensitive individuals (Dong et al., 2008; Feder et al., 1996; Tomanek and Somero, 1999; Welte et al., 1993). But what makes the present study unique is that while the canonical heat shock genes were actively induced in response to heat stress (see below), heat shock gene expression was indistinguishable between tropical and Vermont embryos. Thus, it appears that either thermal adaptation has proceeded via a unique physiological path in fly embryos, or the thermal physiology of embryogenesis is different from other cellular contexts, such that ROS signaling may be a more critical factor than proteostasis. Because ROS signaling mediates various cellular and developmental processes (Petrova et al., 2018; Syal et al., 2020), these broader transcriptomic patterns of differentiation between tropical and Vermont embryos may have resulted from the differential response to oxidative stress. Making the causal link between oxidative stress and the more broadscale patterns of transcription across development would require targeted manipulation of the expression of oxidative stress genes, which is beyond the scope of the present study. Nonetheless, this would be a worthwhile endeavor to pinpoint the mechanism(s) underlying embryonic heat tolerance more precisely.

### A robust early embryonic transcriptomic response

To those who study early embryonic development, perhaps the most surprising result of the present study is that early embryos had any sort of transcriptional response to temperature. Under the current paradigm of metazoan early development, this transcriptional response would not be possible because zygotic transcription is presumed to be silent until the maternal-to-zygotic transition (Tadros and Lipshitz, 2009), which has been shown to take place after 2 hours post-fertilization in *D. melanogaster* (Ashburner et al., 1989; Bashirullah et al., 2003; Bushati et al., 2008; De Renzis et al., 2007; Edgar and Datar, 1996). The embryos in the present study were sampled at 0–1-hour post-fertilization, yet thousands of transcripts exhibited changes in abundance following heat stress, indicating that early embryos are more transcriptionally active than previously described. Some reports demonstrate zygotic transcription in *D. melanogaster* before the MZT, at or prior to nuclear cycle 8 or as early as one hour post fertilization, but these studies show the zygotic expression of very few genes, as embryos were developed under the benign condition of a constant 25°C (Ali-Murthy et al., 2013; Vastenhouw et al., 2019). In contrast, we demonstrate a wide-spread transcriptomic response that is reminiscent of the canonical heat shock response (Gasch et al., 2000; Lindquist, 1986), which involves shifts in gene expression that mitigate protein unfolding (Kültz, 2005; Richter et al., 2010; Somero et al., 2017). Indeed, we observed robust induction of many genes that encode molecular chaperones, including *Hsp70Bb*, *Hsp23*, *Hsp68*, and *Hsp90* (Fig. 6; pink module of WGCNA; Fig. S4). Importantly, most of these genes were highly induced following heat stress, thus demonstrating an active embryonic transcriptional response. We acknowledge that this is not the first observance of heat shock protein expression in early fly embryos (Graziosi et al., 1980), but this is the first demonstration of a transcriptomic response of heat shock genes in early embryos.

Two possible explanations account for this widespread early onset of zygotic transcription upon acute heat shock. First, these results could indicate a shift in the maternal to zygotic transition (MZT) due to the positive relationship between temperature and reaction rates (Somero et al., 2017). Embryos develop faster in hotter temperatures (Kuntz and Eisen, 2014), and we saw increased expression of transcripts involved in developmental processes at hotter temperatures, including pathways in cellular organization and morphogenesis. We also observed that 2,921 transcripts decreased in abundance with increasing temperature. Most of the transcripts that decreased in abundance likely represent maternal transcripts that were being degraded (Atallah and Lott, 2018; Omura and Lott, 2020) because they were present at highest abundances at 25°C. The degradation of these maternal transcripts may be an outcome of the onset of zygotic transcription, which involves the concomitant degradation of maternal mRNAs when the zygotic genome begins to be expressed (Ali-Murthy et al., 2013; De Renzis et al., 2007; Tadros and Lipshitz, 2009).

Alternatively, these transcriptomic responses to increasing temperature could signify a more specific molecular response to heat stress that is distinct from the transcriptomic patterns of the MZT. The most robust changes in gene expression that we observed were among transcripts that encode molecular chaperones, and the induction of heat shock genes does not take place during the MZT (Casas-Vila et al., 2017; Lefebvre and Lécuyer, 2018). Meanwhile, the degradation of thousands of maternal transcripts could be a component of the heat shock response itself, which characteristically involves the downregulation of thousands of genes (Buckley et al., 2006; Gasch et al., 2000; Leemans et al., 2000; Lockwood et al., 2010). In the canonical model of the heat shock response, the expression of molecular chaperones increases, but the expression of thousands of other genes decreases (Somero et al., 2017). This reduction in gene expression serves to halt the production of proteins that could unfold, denature, and cause further risk to the cell (Richter et al., 2010). Thus, our observed transcriptional responses to heat stress suggest a strong heat shock response in early *D. melanogaster* embryos that is distinct from the transcriptional characteristics of the MZT; however, we cannot rule out the possibility of an early shift in the MZT based on these data alone.

## Conclusion

In the context of changing environments, there is much interest in predicting responses to thermal selection (Angilletta, 2009; Pinsky et al., 2019; Sunday et al., 2012). Based on the data we present herein; we predict that early embryonic heat tolerance will respond to future selection because (1) there is standing genetic variation for this trait within and among populations, (2) there is gene flow between low and high latitude populations (Kapun et al., 2021; Pool and Aquadro, 2007; Pool et al., 2012; Pool et al., 2016), and (3) the requisite tweaks to the transcriptome are possible without disruptions to normal developmental processes. It’s important to note that the evolutionary dynamics of adaptation of early embryonic traits may be complex due to interactions between maternal and zygotic genomes. Selection likely acts on both the maternal factors that are loaded into eggs, which do not directly experience the environment of the egg, and the zygotic factors that are expressed in direct response to environmental changes. To our knowledge, the extent to which interactions between maternal and zygotic factors constrain and/or facilitate adaptive responses has not been explored in the context of climate change.

## ACKNOWLEDGEMENTS

We thank Emily Dombrowski, Gwen Ellis, Melissa Pespeni, Sara Helms Cahan, and Seth Frietze for helpful suggestions. This work was funded by NSF grant IOS-1750322 to BLL.

## AUTHOR CONTRIBUTIONS

BLL conceived the study. EEM and BLL designed the experiment. EEM conducted the experiment. EEM, TSO, and BLL analyzed the data. EEM and BLL wrote the manuscript.

## DATA AVAILABILITY

The raw sequence data have been uploaded to the NCBI short read archive (Accession: PRJNA1103862).

