## Supplemental Figures for "Heat tolerance, oxidative stress response tuning, and robust gene activation in early-stage *Drosophila melanogaster* embryos"

### Figure S1

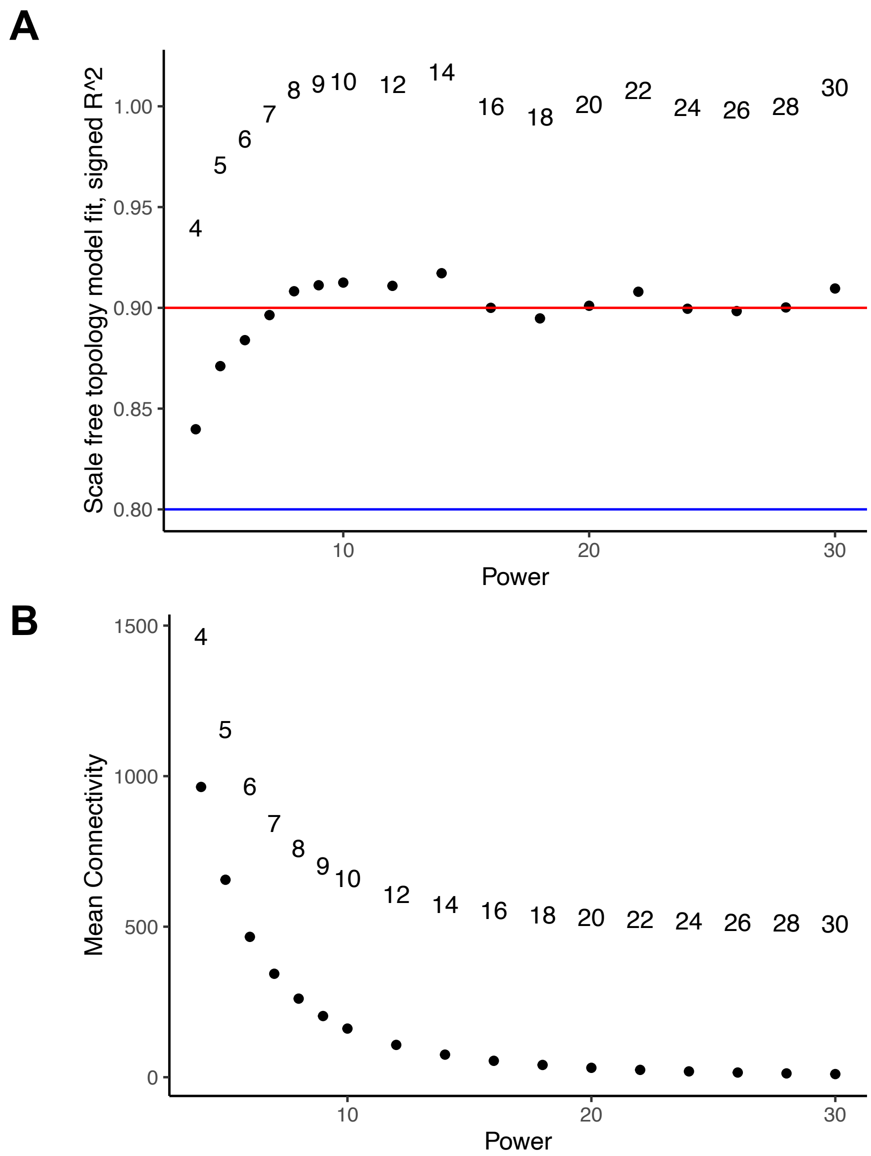


**Figure S1.** WGCNA network properties. (A) Soft power threshold vs. model fit ($R^2$). (B) Soft power threshold vs. mean connectivity.

**Figure S2**

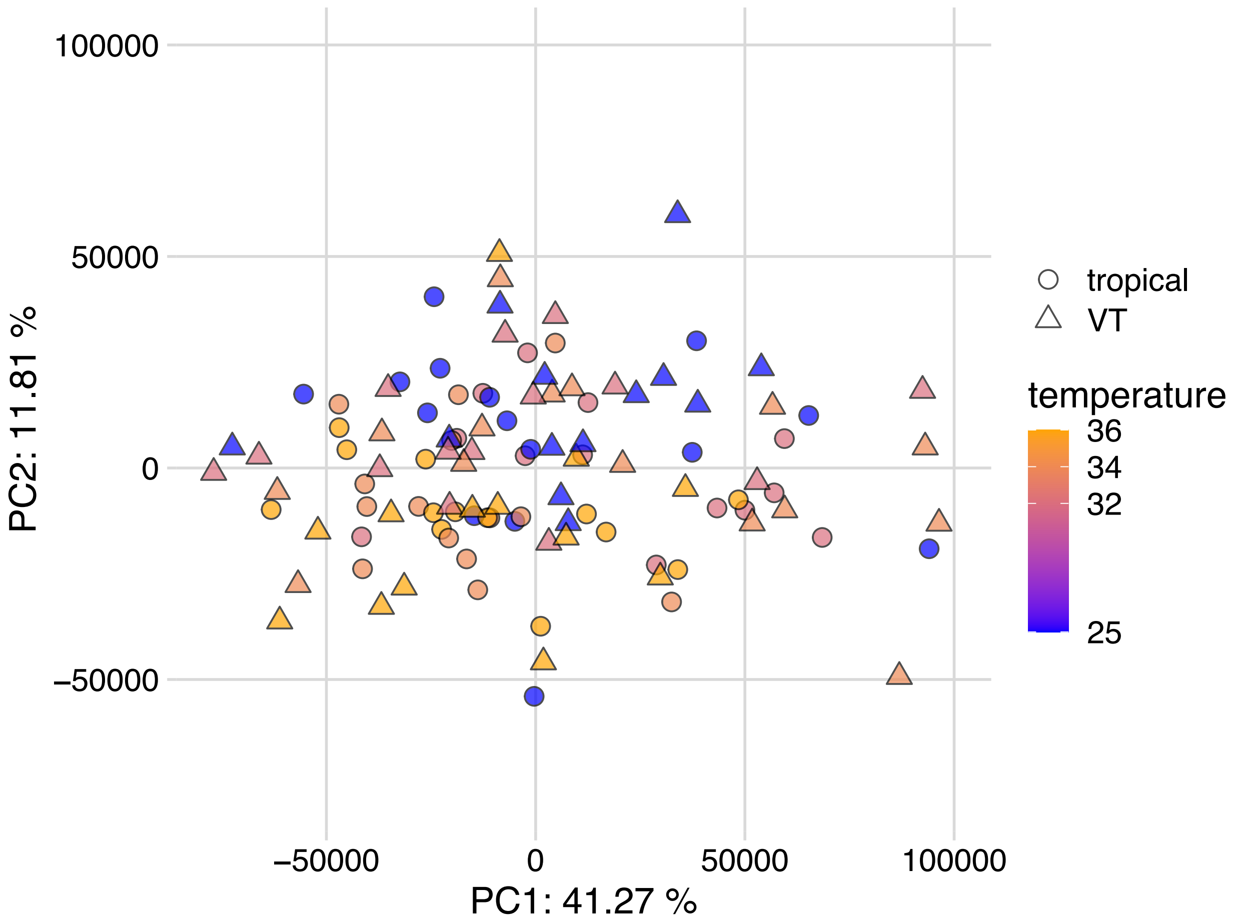


**Figure S2.** Principal component analysis (PCA) of normalized transcript abundance among 110 embryonic samples. The first two principal components together describe 53% of the total variation in transcript abundance. Shape indicates region of origin and color indicates heat shock temperature (°C).

#### Figure S3

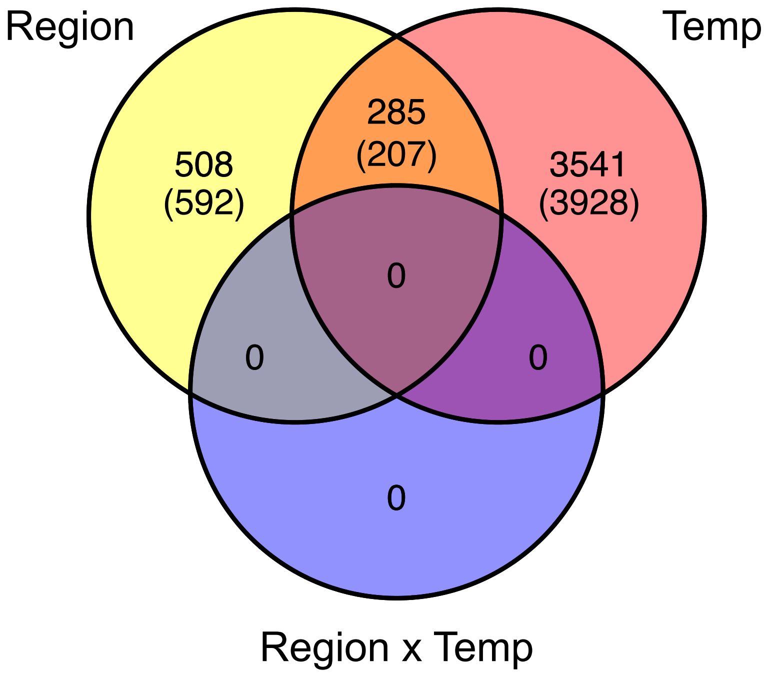


**Figure S3.** Number of significantly differentially expressed genes, with number of transcripts in parentheses, for the main effects of region, temperature or region x temperature interaction (DESeq2; LRT, FDR < 0.05).

### Figure S4

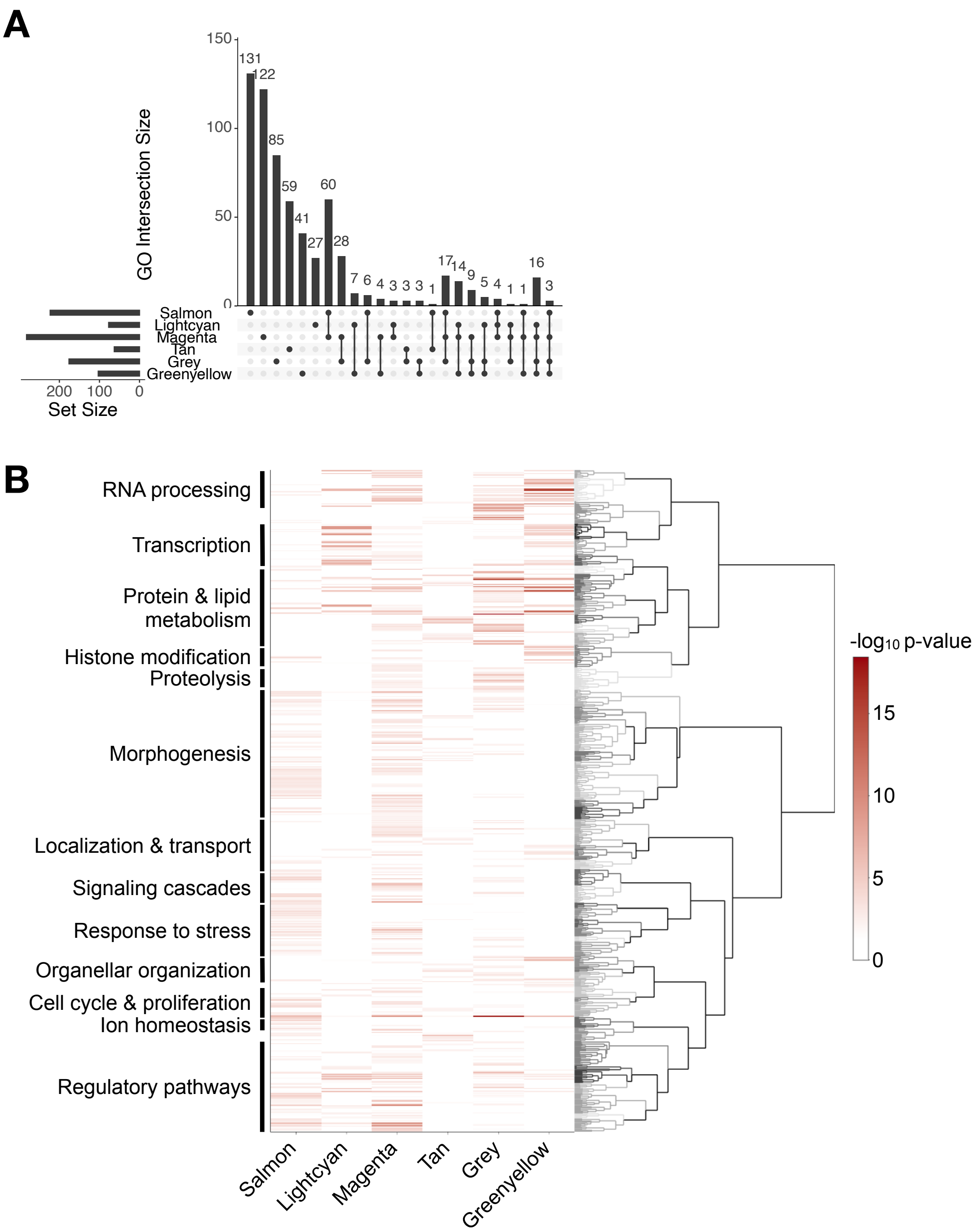


**Figure S4.** Functional enrichment of GO biological process categories among the gene expression modules correlated with embryonic heat tolerance. (A) The number of enriched categories in each module and overlap among modules. (B) The degree to which GO categories were enriched, with the darker color indicating greater enrichment. The dendrogram indicates the relationships among GO categories, with functionally related categories clustered together on the tree. GO categories are further summarized by broad functional classifications by the authors, based on FlyBase (release FB2023_06) (Jenkins et al., 2022). Note that some gene modules exhibit enrichment of a broad array of functional categories while other modules are more specific to particular functional categories.

### Figure S5

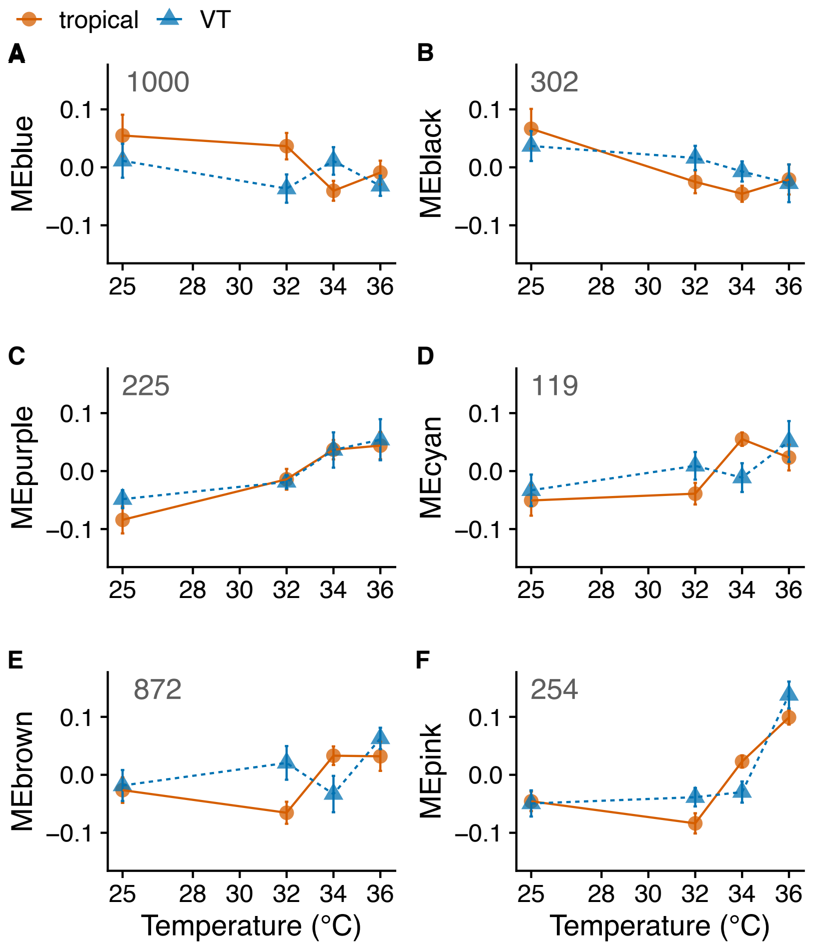


**Figure S5.** ME values across temperature for the six modules with significant correlations to temperature. Note that ME reflects the pattern of expression common to all genes in a particular module, with higher values indicating greater abundance of RNA. Numbers in each panel indicate the number of genes in that module.
